# End-to-end evaluation of white matter microstructure of the visual pathway in asymmetric glaucoma

**DOI:** 10.1101/2025.10.20.683506

**Authors:** Daniela Coutiño, Judith Guerrero, César Arturo Domínguez-Frausto, Marlene Garcia-Guillén, Ricardo Coronado-Leija, Alonso Ramírez-Manzanares, Erick Hernández-Gutiérrez, Maxime Descoteaux, Martin Ayala, Mariana Badillo, Luis Concha

## Abstract

Diffusion magnetic resonance imaging is a non-invasive neuroimaging technique that enables in vivo evaluation of white matter microstructure, providing sensitivity to tissue abnormalities caused by disease. Glaucoma, the second leading cause of blindness worldwide, is characterized by progressive loss of retinal ganglion cells and axonal damage in the optic nerve, leading to degeneration along the entire visual pathway. This degeneration includes secondary effects on fiber crossings within the optic chiasm, which are challenging to characterize with conventional diffusion models. In this study, we evaluated 31 patients with asymmetric glaucoma and 31 healthy controls using advanced diffusion magnetic resonance imaging methods, including Diffusion Tensor Imaging, Constrained Spherical Deconvolution, multi-tensor fit via Multi-Resolution Discrete Search method, and Fixel-Based Analysis. We found significant differences of diffusion metrics in white matter tracts of the visual system, including the optic nerve, optic chiasm, optic tracts, and optic radiations. Moreover, diffusion metrics correlated with clinical ophthalmological parameters such as cup-to-disc ratio, visual field mean deviation, and retinal nerve fiber layer thickness. These findings support the use of advanced diffusion magnetic resonance imaging models as sensitive tools for detecting Wallerian degeneration and resolving complex white matter architecture in the human visual pathway, and demonstrate their utility to study other fiber-crossing regions throughout the brain.

## Introduction

Diffusion-weighted magnetic resonance imaging (dMRI) enables noninvasive assessment of tissue microstructure [1–3]. Diffusion tensor imaging (DTI) [4] is the most commonly used method, given its efficiency and wide availability. However, DTI models diffusion in only one prominent direction, rendering it unsuitable for regions with complex fiber configurations, such as fiber crossings—which are present in the majority of human white matter [5–7]. This limitation causes the inaccurate estimation of oblate or spherical tensors in regions containing multiple fiber orientations, thereby reducing fractional anisotropy (FA). Moreover, in cases when one of the crossing fiber populations is degenerated, the remaining bundle dominates the diffusion signa, and the resulting tensor appears to be anisotropic (i.e., with increased FA), leading to incorrect biological interpretations [8].

Other methods for analyzing the diffusion signal can resolve multiple fiber orientations. Independent estimation of diffusion tensors for each fiber bundle within each voxel can be achieved with Multi-Resolution Discrete Search (MRDS), a method with relatively low data acquisition requirements and robustness to noise [9]. Similarly, Constrained Spherical Deconvolution (CSD) [10] can identify one or more fiber elements (fixels) and derive apparent fiber density (AFD) values independently for each fiber bundle [11]. In previous work using an animal model of unilateral retinal ischemia and high-resolution *ex vivo* dMRI, our group showed that multi-tensor approaches, as well as CSD analyses, are able to correctly identify a known region of fiber crossings in the optic chiasm and, importantly, provide independent and accurate microstructural information related to the degenerated and intact axonal populations that cross within this structure [12]. Moreover, the per-bundle diffusion metrics derived from MRDS and CSD were tightly correlated with the degree of degeneration of the optic nerves assessed with electron microscopy. The ability to disentangle intact and degenerated fiber populations through bundle-wise analyses has immediate applications to characterize other regions of white matter with complex configurations [13], yet they require further validation when derived from dMRI acquisitions with parameters constrained to clinical settings (i.e., relatively short acquisition times and lower resolution).

In this work, asymmetric glaucoma, characterized by unequal severity of damage between the two eyes, serves as a case study that allows for the spatial assessment of diffusion metrics throughout the visual pathway. Glaucoma is a leading cause of irreversible blindness worldwide that affects 3.54% of the population aged 40 to 80 years [14]. It causes progressive degeneration of the retinal ganglion cells and leads to gradual peripheral vision loss [15]. Generally, by the time of diagnosis, 30–50% of retinal ganglion cells have already been lost. The current evaluation of patients with glaucoma relies on visual field parameters such as mean deviation (VF-MD) and and on optical coherence tomography (OCT) metrics, including the vertical cup-to-disc ratio (vCD) and retinal nerve fiber layer (RNFL) thickness, which reflect the degree of retinal degeneration. The axons that extend from retinal ganglion cells form the optic nerve, and suffer Wallerian degeneration upon damage to their soma. Several studies have shown correlations between diffusion metrics of the optic pathway and said clinical parameters [16–19]. However, previous reports have avoided analyzing the optic chiasm due to its complex crossing fibers.

This study investigates the potential of advanced dMRI to detect *in vivo* structural alterations of the optic pathway in patients with asymmetric glaucoma. We focus on the optic chiasm as a model of crossing-fiber architecture, and the asymmetric pattern of retinal damage caused by glaucoma provides a clear scenario in which two fiber populations with differing degrees of degeneration cross in a known anatomical structure. Our objectives are (*i*) to validate per-bundle dMRI analyses in clinically-accessible dMRI data, with the goal that such validation may extend to other white-matter regions with crossing fibers throughout the brain and (*ii*) to characterize white matter damage throughouth the entire visual pathway and assess the correlation of diffusion metrics with clinical parameters.

## Materials and methods

### Participants

We recruited adult patients with asymmetric glaucoma from the Glaucoma Service at the Mexican Institute of Ophthalmology (IMO). A total of 1,926 clinical records of patients evaluated in the last 12 months were screened. The majority of patients were excluded due to symmetric glaucoma (802), age outside the inclusion range (654), single functioning eye (158) incomplete diagnostic studies (94), or other clinical and radiological contraindications, such as the presence of pacemakers, metallic implants, or mobility aids incompatible with MRI (12). Despite potential eligibility, 171 patients were not willing to participate or were not found based on their contact information. After applying these criteria, 35 patients underwent imaging. Four patients were excluded due to unexpected radiological findings. This resulted in a final cohort of 31 patients for analysis (14 females and 17 males) with a mean age of 59 ± 14 years old. Patients were considered to have clearly asymmetric glaucoma when there was an interocular difference of their vertical cup-to-disc ratio (vCD) ≥ 0.2, together with an interocular difference in retinal nerve fiber layer (RNFL) thickness ≥ 10 µm on OCT [20, 21]. Thirty-one age-matched healthy controls (15 females and 16 males) were also included. Participation in this study was voluntary and independent from treatment. All procedures were conducted in accordance with the Declaration of Helsinki. The protocol was approved by the Research Ethics Committee of the Institute of Neurobiology and by the Bioethics Committee of the IMO.

### Imaging

The dataset was acquired on a 3 T Discovery MR750 scanner (GE Healthcare, Milwaukee, WI, USA) with a 32-channel head coil. We acquired multi-shell dMRI using two different methods, both had three non-zero b-shells (b = 800, 1500, 2500 s/mm^2^) with diffusion sensitization in 16, 32, and 64 different directions. Six b=0 s/mm^2^ volumes were also acquired. A Multiplexed sensitivity-encoding (MUSE) DWI sequence [22] was used to acquire an axial slab encompassing the visual pathway and oriented parallel to the optic nerves (3 shots, 28 slices, resolution of 1.7×1.5×1.7 mm^3^ interpolated by the scanner to 0.86×0.86 in-plane resolution, TR=3500 ms, TE=80 ms). This multi-shot scan strategy allowed for the reduction of geometric distortions that occur at the base of the skull and affect the optic nerves, chiasm, and tracts. Whole-brain dMRI were acquired with a simultaneous multi-slice sequence (multiband factor=2, TR=4000 ms, TE=80 ms, isotropic resolution of 1.7 mm). Additionally T1-weighted images (0.8 mm isotropic resolution) and T2-weighted images (1 mm isotropic resolution) were acquired for screening purposes. Total acquisition time was approximately 40 minutes.

### Preprocessing

All dMRI data sets were first denoised using either Marchenko–Pastur principal component analysis (MP-PCA) [23, 24] for the whole-brain data sets, or Patch2Self [25] for the in-plane interpolated MUSE-DWI. Gibbs ringing artifacts were removed using the subvoxel-shifts method [26]. Subject motion, as well as magnetic susceptibility and eddy-current induced distortions, were corrected with topup and Eddy tools, available in the FSL toolbox (version 6.0.7.4) [27]. Bias field inhomogeneities were then corrected using the N4 algorithm [28].

### Voxel-wise analysis of dMRI

Diffusion tensor and kurtosis imaging (DTI and DKI, respectively) were estimated using DIPY [29, 30]. The diffusion tensor yielded parameters of fractional anisotropy (FA) and mean diffusivity (MD). The diffusion kurtosis measures the non-Gaussianity of water diffusion and provides additional information on tissue complexity [31]. Mean kurtosis (MK), axial kurtosis (AK), and radial kurtosis (RK) were calculated at each voxel. Regions of interest (ROI) were manually drawn outlining each optic nerve using mrview (MRtrix 3.0.4) [32], and average DTI and DKI metrics were obtained for each ROI.

### Per-bundle analysis of dMRI

#### Constrained Spherical Deconvolution (CSD)

This method was used to calculate the Apparent Fiber Density (AFD) [11] from the MUSE DWI acquisition, which provided finer detail and minimized geometric distortions around the optic chiasm and nerves. White matter response functions were estimated using a recursive calibration [33], applied within a whole-brain mask. As this analysis was restricted to the optic nerves and chiasm, single-tissue CSD was applied to obtain the white matter fiber orientation distributions (FODs). The resulting FODs were normalized to allow inter-subject comparison. These normalized FODs were used to generate tractography, encompassing fibers from both optic nerves, the optic chiasm, and the initial portion of the optic tracts, as shown in Fig 1. FODs were converted into fixels, restricting the maximum number to two fixels per voxel (i.e., expecting one fixel at the level of optic nerves and tracts, and two fixels within the chiasm), and fixel-wise AFD maps were derived within manually-defined ROIs in the optic nerves and chiasm

**Fig 1.**
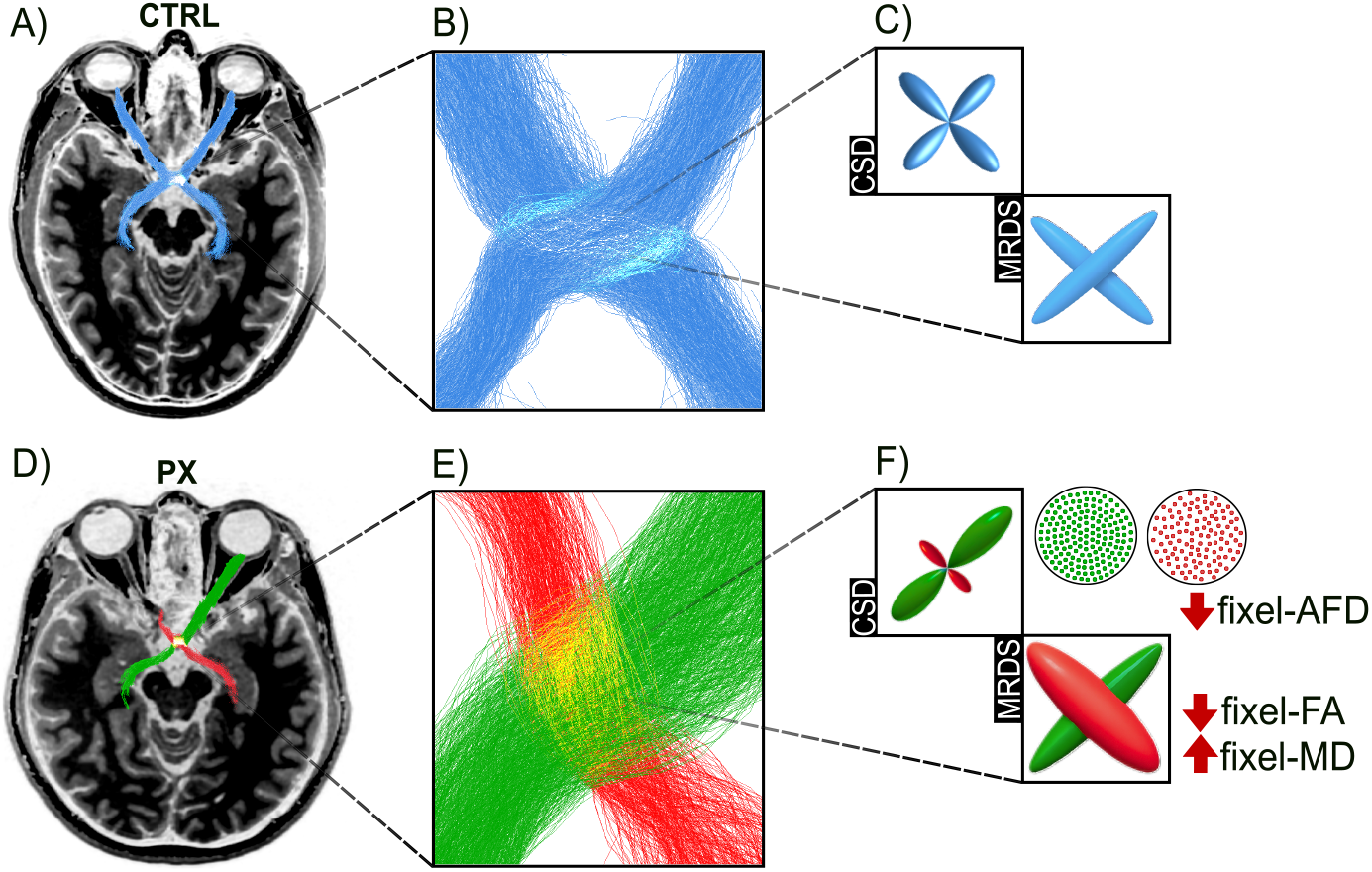
Tractography and bundle-specific diffusion metrics in the optic chiasm of a control subject (A–C) and a patient with asymmetric glaucoma (D–F). Panels A and D show whole-brain T1-weighted images with tractography of the visual system overlaid. Panels B and E display zoomed views of the optic chiasm, highlighting the crossing of nasal (contralateral, green) and temporal (ipsilateral, red) fibres. Panels C and F provide schematic representations of a voxel at the fibre crossing. In the control subject, both multi-resolution discrete search (MRDS) and constrained spherical deconvolution (CSD) depict symmetric fiber populations. In the patient, MRDS shows reduced fixel-FA and increased fixel-MD in the ipsilateral fibers (red colored fixels), while CSD demonstrates reduced apparent fiber density (AFD). These alterations illustrate the sensitivity of bundle-specific metrics to detect glaucoma-related degeneration in regions with complex fiber architecture.

#### Multi-Resolution Discrete Search (MRDS)

A region of interest (ROI) was manually drawn to delineate each optic chiasm, where one or two diffusion tensors were fit at each voxel, then used to derive maps of fixel-FA and fixel-MD corresponding to the ipsilateral and contralateral sides of the eye affected by glaucoma. Although MRDS includes different model selectors to determine the number of tensors, in this study we employed the Track Orientation Density Imaging (TODI) method as the selection strategy [34]. This approach uses tractography obtained with CSD to generate voxel-wise track orientation distributions, which provides a more consistent representation of fiber orientation than FODs [35]. The TODI was used to derive a Number of Fiber Orientations (NuFO) map, which was constrained to ≤ 2, corresponding to the two main fiber populations in the chiasm (one from each optic nerve) [36]. Next, the diffusion tensors best aligned with each optic nerve (either ipsilateral or contralateral to the most affected eye) were selected, from which we derived FA and MD (referred to as fixel-FA and fixel-MD, respectively, to disambiguate these from the standard DTI metrics obtained in the optic nerves). The two metrics were separately averaged for each orientation (i.e., ipsilateral and contralateral to the most affected eye).

#### Fixel-based analysis

The DWI obtained with simultaneous multi-slice acquisition providing full brain coverage were resampled to 1.25 mm isotropic resolution after pre-processing. Response functions for white matter, gray matter, and cerebrospinal fluid were estimated using the unsupervised approach described by Dhollander [37, 38], followed by multi-tissue CSD (MSMT-CSD) to compute FODs [39]. Individual FOD images were normalized and a population FOD template was generated using nonlinear registration [40], to which all subjects were registered. From this template, fixel-wise metrics of Fiber Density (FD), Fiber Cross-section (log-FC), and Fiber Density and Cross-section (FDC) were computed [41]. A whole-brain tractogram of 20 million streamlines was generated from the template and filtered to 2 million streamlines using SIFT to reduce reconstruction biases [42]. Fixel-wise metrics were smoothed with a 10-mm FWHM Gaussian kernel, and statistical analysis was performed using the connectivity-based fixel enhancement (CFE) approach with 5000 permutations, comparing group means with age as a covariate [43].

### Statistical analyses

Data normality was assessed with the Shapiro–Wilk test for each eye (right and left in controls; contralateral and ipsilateral to the most affected eye in patients). Depending on distribution, paired Student’s t test or Mann–Whitney test was used to compare right vs. left optic nerves in controls. As no significant differences were found, an average per control subject was calculated. Average values of the optic nerves of the control group were then separately compared to metrics derived from the patients’ optic nerves according to their relation to the most affected eye using unpaired Student’s t or Mann–Whitney tests. Finally, paired comparisons were performed between the two optic nerves in patients. Statistical significance was considered as p*<*0.05.

### Correlations between diffusion metrics and clinical parameters

Bivariate correlations were performed to explore the relationship between diffusion metrics and clinical parameters of glaucoma severity. These correlations are summarized in the matrix shown in Fig 5, where each cell with a dot indicates a significant correlation and the correlation coefficient (*r*) is color-coded. Additional correlations between diffusion metrics are provided in Supplementary Fig 1.

### Data availability

Data will be made available upon article publication.

## Results

In patients with asymmetric glaucoma, the nerve related to the most affected eye (ipsilateral) showed significantly altered values compared to the contralateral eye (Fig 2). Nearly all voxel-wise metrics showed clear asymmetry between the two optic nerves of patients, characterized by reduced FA, AFD, MK, RK, as well as increased MD, in the optic nerves ipsilateral to the most affected eye. The contralateral optic nerves of patients showed statistical difference to control subjects in FA and AFD.

**Fig 2.**
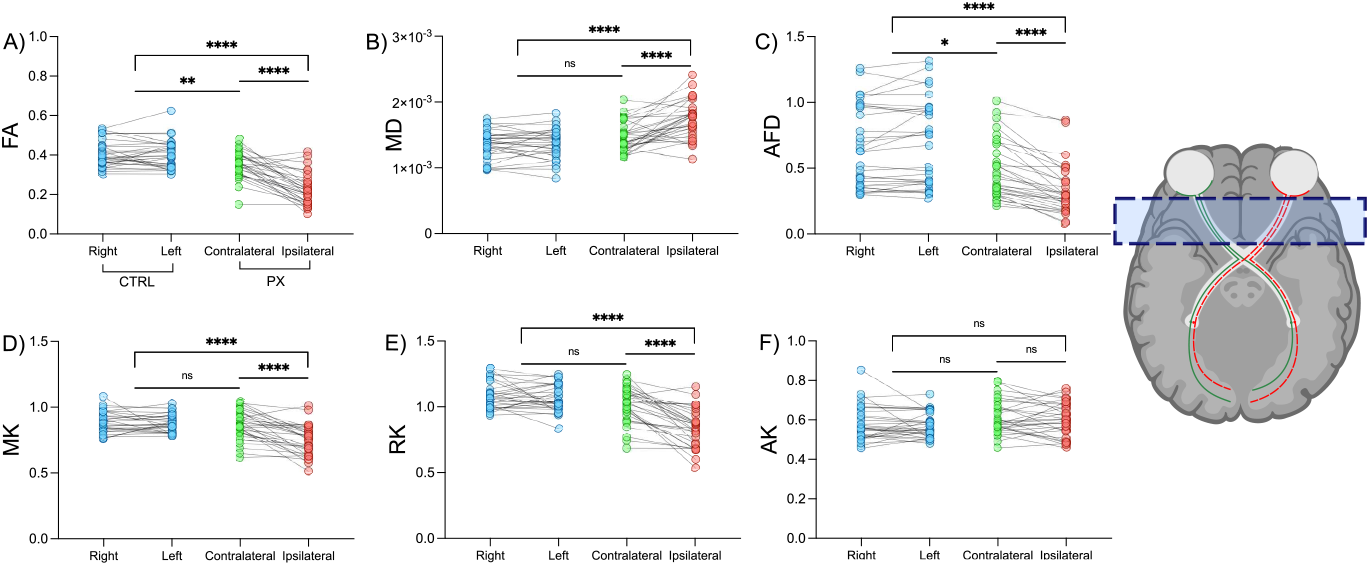
Comparisons of diffusion metrics in the optic nerves between healthy controls and patients with asymmetric glaucoma. Diffusion metrics (FA, MD, AFD, MK, RK, and AK) are shown separately for fibers stemming from right or left eyes in controls (blue), and from the contralateral (green) or most affected/ipsilateral eyes (red) in patients. Voxel-wise analysis was performed at the level of the optic nerves (A–F). ns = not significant; *p *<*0.05, **p *<*0.01, ***p *<*0.001, ****p *<*0.0001.

Per-bundle analyses of the optic chiasm showed significant reductions of fixel-FA (MRDS) between the two fiber populations (i.e., stemming from ipsilateral and contralateral eyes), as well as between the affected eye and the control group, but not between the control group and the contralateral eye, consistent with findings from AFD using CSD (Fig 3). For fixel-MD, a significant increase was identified both intragroup (most affected vs. contralateral eye) and intergroup (patients vs. controls).

**Fig 3.**
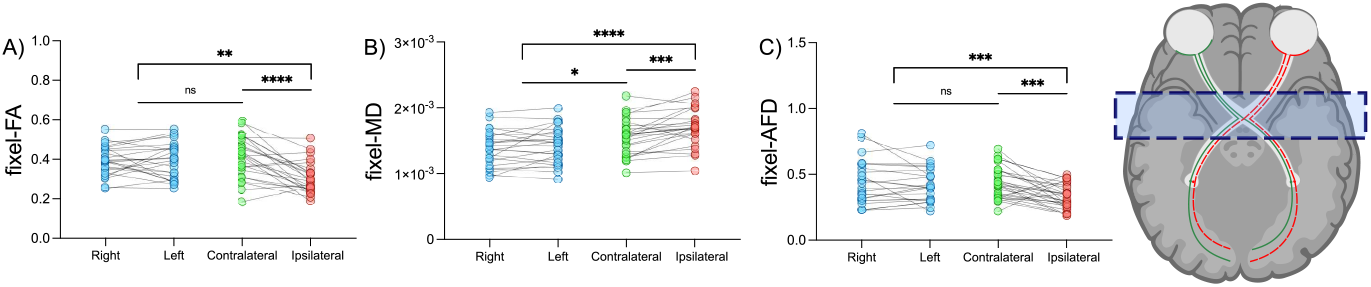
Comparisons of diffusion metrics in the optic chiasm between healthy controls and patients with asymmetric glaucoma. Fixel-wise metrics (fixel-FA, fixel-MD, and fixel-AFD) are shown separately for fibers stemming from right or left eyes in controls (blue), and from the most affected (ipsilateral, red) or contralateral (green) eyes in patients. Blue shading in the anatomical illustration indicates the analysis level at the optic chiasm. ns = not significant; *p *<*0.05, **p *<*0.01, ***p *<*0.001, ****p *<*0.0001.

Fixel-based analysis revealed significant reductions in patients with asymmetric glaucoma in fiber density (FD), fiber cross-section (Log-FC), and the combined metric FDC in both the optic tract and optic radiations compared to controls (Fig 4). These results reflect microstructural alterations (FD), macrostructural changes (Log-FC), and combined effects (FDC). Only reductions of FD within the optic tracts and of FDC within both optic tracts and optic radiations were statistically significant after cluster-based correction for multiple comparisons.

**Fig 4.**
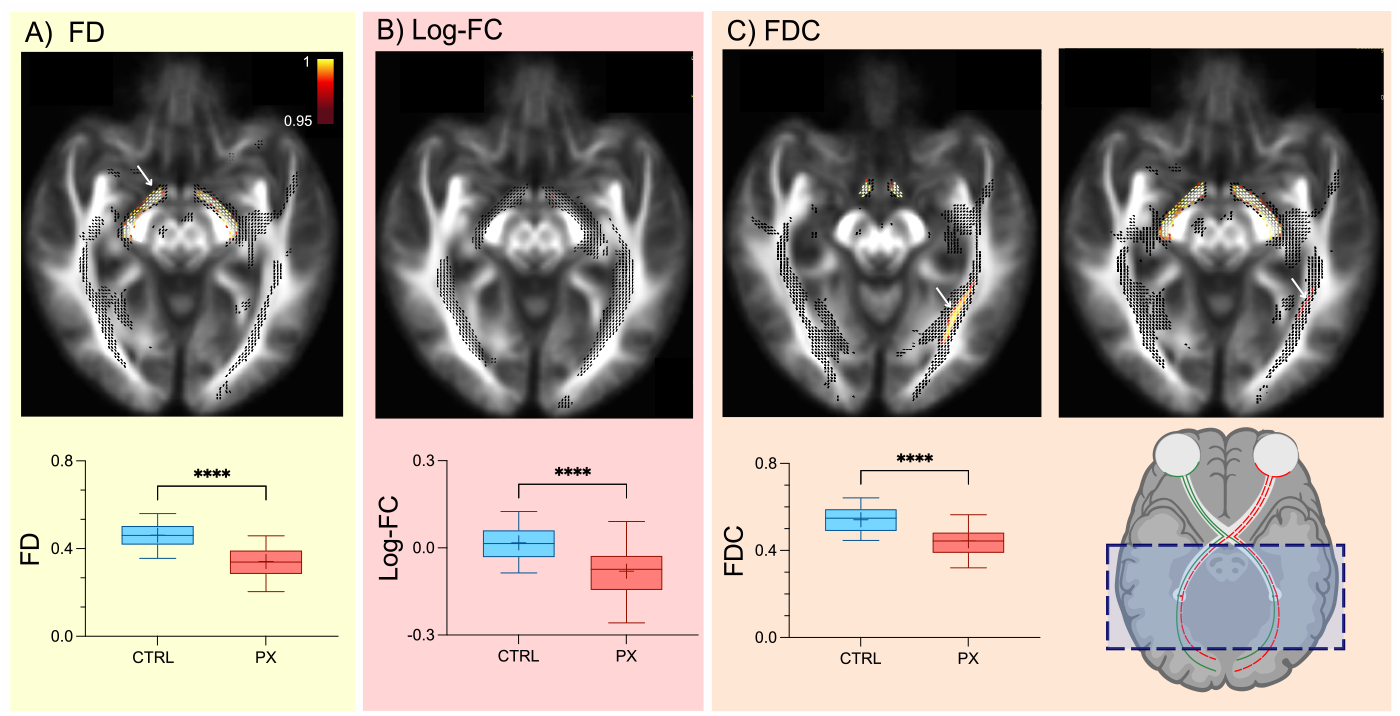
Fixel-based group comparisons in the optic tract and optic radiations. A) Fibre Density (FD), B) Fibre Cross-section (log-FC), and C) Fibre Density and Cross-section (FDC) showed significant reductions in patients with asymmetric glaucoma (red) compared to healthy controls (blue). Top panels display statistical fixel maps overlaid on the population template. Fixels are visualized (black lines) if statistically significant (uncorrected p*<*0.05), and color-coded (1-p) if significant after correction using connectivity-based fixel enhancement (CFE). Lower panels show group boxplots for each metric at the regions indicated by the arrows in the top panels. ****: p*<*0.0001.

Significant correlations were observed between diffusion MRI metrics and clinical parameters along the white matter of the visual pathway (Fig 5). As expected in an asymmetric glaucoma cohort, the ipsilateral eye showed stronger associations with clinical measures, such as FA correlating negatively with vCD and fixel-FA correlating positively with VF-MD. These examples illustrate the direct link between microstructural degeneration and clinical severity. In contrast, contralateral eyes exhibited a spectrum of damage, reflecting that in some patients the asymmetry was mainly due to differences in the degree of impairment between eyes. Consequently, correlations with clinical parameters were less consistent on the contralateral side; these results are presented in Supplementary Fig. S1. Additional correlations between diffusion metrics themselves are also provided in Supplementary Fig S1.

**Fig 5.**
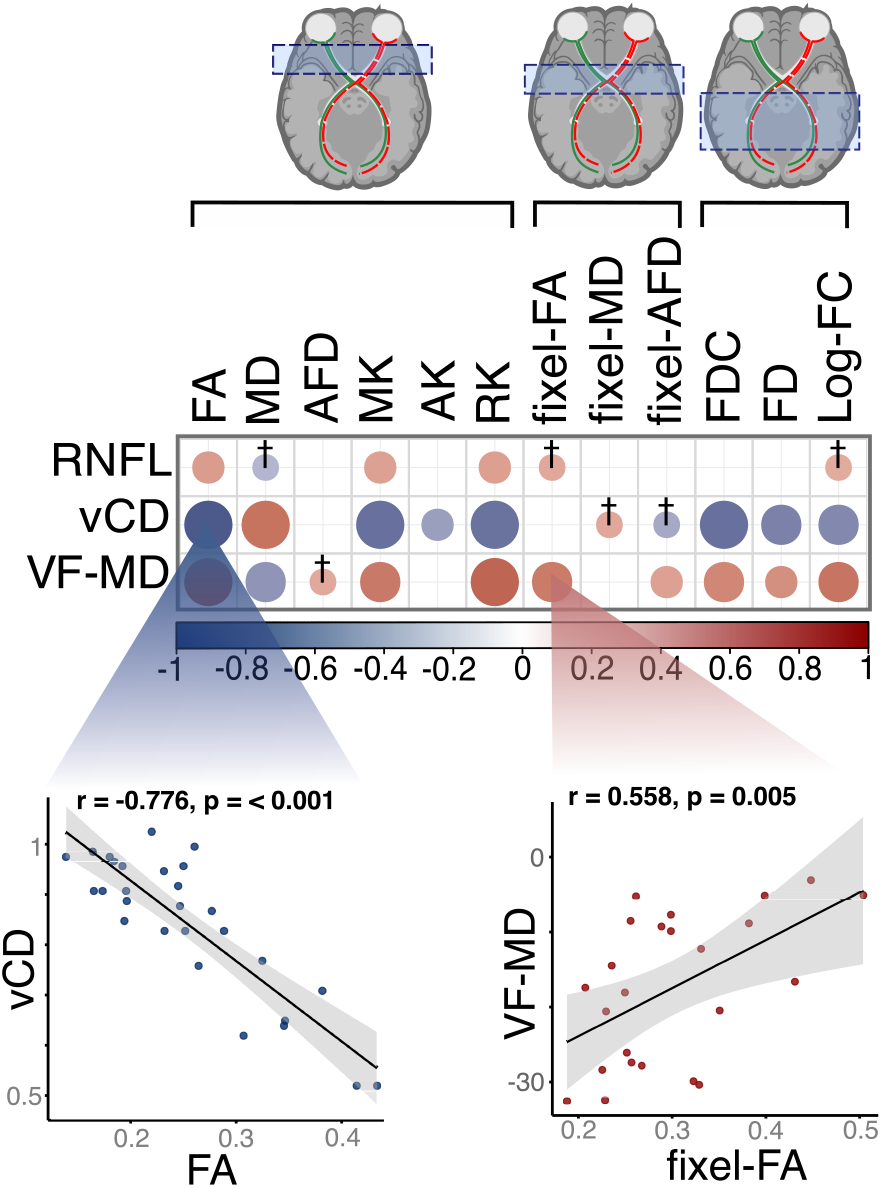
Correlation matrix between diffusion MRI metrics and clinical parameters. Significant correlations are shown as circles (blue = negative, red = positive). Circle diameters reflect the significance of the correlation (p*<*0.05), and crosses indicate trends (0.05*<*p*<*0.1). Scatter plots illustrate two exemplary associations. Clinical parameters and diffusion metrics of the optic nerve and chiasm correspond to the most affected eye; the average diffusion metrics of both hemispheres is used for the optic radiations.

## Discussion

This study provides evidence of white matter damage throughout the entire visual system in patients with asymmetric glaucoma. We show that different clinically-feasible diffusion metrics provide complementary information regarding the microarchitecture of the optic nerves, tracts, chiasm, and radiations. Moreover, these metrics correlate with disease severity, providing insights into the down-stream degeneration of brain tissue secondary to glaucoma and, importantly, highlighting their potential as monitoring biomarkers.

The sensitivity of DTI for the detection of Wallerian degeneration in the optic nerve is well-known. Indeed, evaluation of the optic nerves of rodents after experimental retinal ischemia served as a test-bed for this purpose in the early days of DTI [44, 45]. These and other evaluations identified the link between axonal injury with axial diffusivity, and axonal density and myelin abnormalities with radial diffusivity [2, 46]. It follows that alterations of DTI metrics that indicate degeneration of the optic nerves have been demonstrated in patients with glaucoma [16–19]. Such findings are replicated in this work, yet, as compared with previous reports of patients with bilateral glaucoma, the population studied here allowed for the association of white matter damage to either the most affected or contralateral eyes. The severity of downstream degeneration of the optic nerves assessed with voxel-wise metrics derived from DTI and DKI matches the asymmetry of glaucoma (Fig 2). Furthermore, there is a tight correlation between diffusion metrics of the optic nerves and the degree of retinal alterations observed in OCT and visual field deviations (Fig 5). These results confirm the sensitivity of diffusion metrics to identify white matter degeneration in simple fiber configurations [47–49].

The optic nerves converge at the level of the optic chiasm, where around half of the axons (from the nasal hemisphere of each retina) cross the midline and interdigitate with axons stemming from the other eye. A major limitation of DTI is its inability to resolve crossing fibers, whereby fractional anisotropy is artifactually reduced in voxels with more than one fiber populations. This has hampered the application of DTI to study the optic chiasm in patients with glaucoma. The chiasm is a very discrete anatomical structure with a complex fiber configuration that can be inferred *a priori*. It therefore serves as an ideal region to provide evidence of the utility of advanced analyses of the diffusion signal. This is not only relevant to the study of the optic pathway, but for the investigation of any region of the brain with crossing fibers, which can account to up to two thirds of the human white matter [7]. Several methods have been proposed to resolve individual orientations of fiber populations that cross within a voxel [50], although not all possess the ability to derive per-bundle diffusion metrics [51]. In this work, we focus on two methods that reliably and independently characterize diffusion for each fiber population, with both methods having relatively lenient acquisition requirements that can be obtained with clinical scanners. The asymmetric presentation of glaucoma in the patients studied here provides a unique opportunity to assess the ability of these methods to detect per-bundle axonal alterations. We show that in the chiasm the multi-tensor fit performed with MRDS shows asymmetry of fixel-FA akin to that seen between the two optic nerves, where the fixels associated to the most affected eye had reduced fixel-FA (Fig 3A). Albeit to a lesser degree, there was also asymmetry of fixel-MD of the two fiber systems in the chiasm of patients, with a slight increase of the fibers related to the most affected eye (Fig 3C). Similar to the multi-tensor fit, CSD was also able to identify the most affected fiber population within the chiasm, which showed marked reductions of AFD (Fig 3C). Also, fixel metrics derived from the multi-tensor fit and CSD (fixel-FA and fixel-AFD, respectively) were positively correlated with visual field deviations (Fig 5). These results in the chiasm of patients with asymmetric glaucoma are in line with our previous exploration of bundle-wise diffusion metrics in the chiasm of rodents with unilateral retinal ischemia, which directly and independently correlate with the number of axons in the corresponding optic nerves evaluated with quantitative histology [12]. In the human condition described here, the thickness of the RNFL and the vCD serve as indicators of retinal damage and act as proxies for the degree of axonal degeneration. However, neither of these metrics showed significant correlations with fixel-wise diffusion metrics in the chiasm. The lack of correlations between retinal parameters and fixel-wise diffusion metrics is likely due to anatomical differences: while nearly all axons decussate at the level of the chiasm in rodents, only 50% cross the midline in humans [52]. The consistency between imaging metrics and clinical severity strengthens the interpretation of our findings. Our current findings show that the ability to separate the diffusion signal into individual fiber bundles allows for a more precise characterization of microstructural changes in regions with crossing fibers. Importantly, these methods are applicable to other regions of the brain with crossing fibers and a wide range of neurodegenerative conditions.

White matter damage secondary to glaucoma is not restricted to the optic nerves, as trans-synaptic alterations have been reported in the optic radiations, and ultimately lead to reduced cortical thickness of the visual cortex (for a recent survey, see [53]). These findings indicate that, although originating in the eye, glaucoma is a condition with complex ramifications throughout the entire visual pathway. Whole-brain FBA clearly identified alterations of the optic radiations in patients with asymmetric glaucoma. Reductions of FD and FDC were observed bilaterally, as expected given the deccussation of fibers in the optic chiasm. The three fixel-wise metrics of the optic radiation significantly correlated with vCD and VF-MD, demostrating a tight link between clinical features and white matter abnormalities even in the most posterior aspects of the visual pathway.

Our findings are in agreement with previous reports that have shown alterations of diffusion metrics of the optic radiations in patients with glaucoma [54]. Voxel-wise analyses, most commonly based on DTI, DKI, or NODDI, have consistently reported reduced FA and increased MD and RD in the optic nerves, optic tracts, and optic radiations, often accompanied by decreases in axial diffusivity or kurtosis-based parameters [55–60]. These alterations correlate with clinical severity, including visual field mean deviation, vertical cup-to-disc ratio, and RNFL thickness. Notably, voxel-wise methods demonstrated that diffusion changes extend beyond the anterior visual pathway into the optic radiations and even extrastriate cortical regions. By contrast, the relatively fewer fixel-wise studies have provided bundle-specific evidence of degeneration. Using FBA, Haykal et al. reported significant reductions of FD, FC, and FDC in different segments of the optic nerve, showing correlations with clinical markers of glaucoma severity [58,61]. These findings highlight that fixel-wise metrics capture both microscopic loss of axonal density and macroscopic reductions in bundle cross-section, offering complementary sensitivity compared to voxel-wise indices.

Interestingly, the most direct assessment of retinal degeneration, namely RNFL, was correlated with diffusion metrics (FA and MK) in the optic nerves, but not in the rest of the visual pathway. For each patient we considered the average RFNL for each eye, which is an over-simplification of the retinal changes seen in glaucoma, that tend to be zonal rather than distributed. Although it is known that the entire visual pathway maintains a retinotopic distribution of fibers and one could theoretically match retinal quadrants with specific regions of white matter, the dMRI resolution achieved herein is insufficient to perform such a detailed analysis. The ability to distinguish two different fiber populations that cross within a voxel is proportional to the angle between them. With the current radial resolution of the diffusion signal used here, both MRDS and CSD should be able to disentangle two fiber populations separated by at least 35-45°. However, higher-resolution dMRI acquisitions have been shown to increase the detectability of fiber crossings and complex configurations [62]. Regardless, it is possible that the two methods may have identified only one fixel in an unknown portion of fixels within the chiasm, where in fact two fiber populations coexist. While conceptually and etymologically the chiasm resembles the greek letter *χ*, its morphology in humans can also resemble the letter H, with axons crossing the midline along the horizontal portion with antiparallel orientations. However, in coronal sections, the axons from either eye meet at an angle given their superior-inferior trajectories [63], thus improving their detection as two distinct fixels.

## Conclusion

Diffusion metrics identified white matter abnormalities throughout the visual pathway of patients with glaucoma indicative of Wallerian and trans-synaptic degeneration, and are useful imaging biomarkers for severity of the disease. Our results provide evidence of the ability of advanced dMRI methods to robustly evaluate white matter characteristics in white matter, even in regions with fiber crossings, where each fiber population can be assessed independently. By extension, our results allow for better interpretation of fixel-wise analyses of white matter beyond the optic pathway.

## Supporting information

**S1 Fig.**
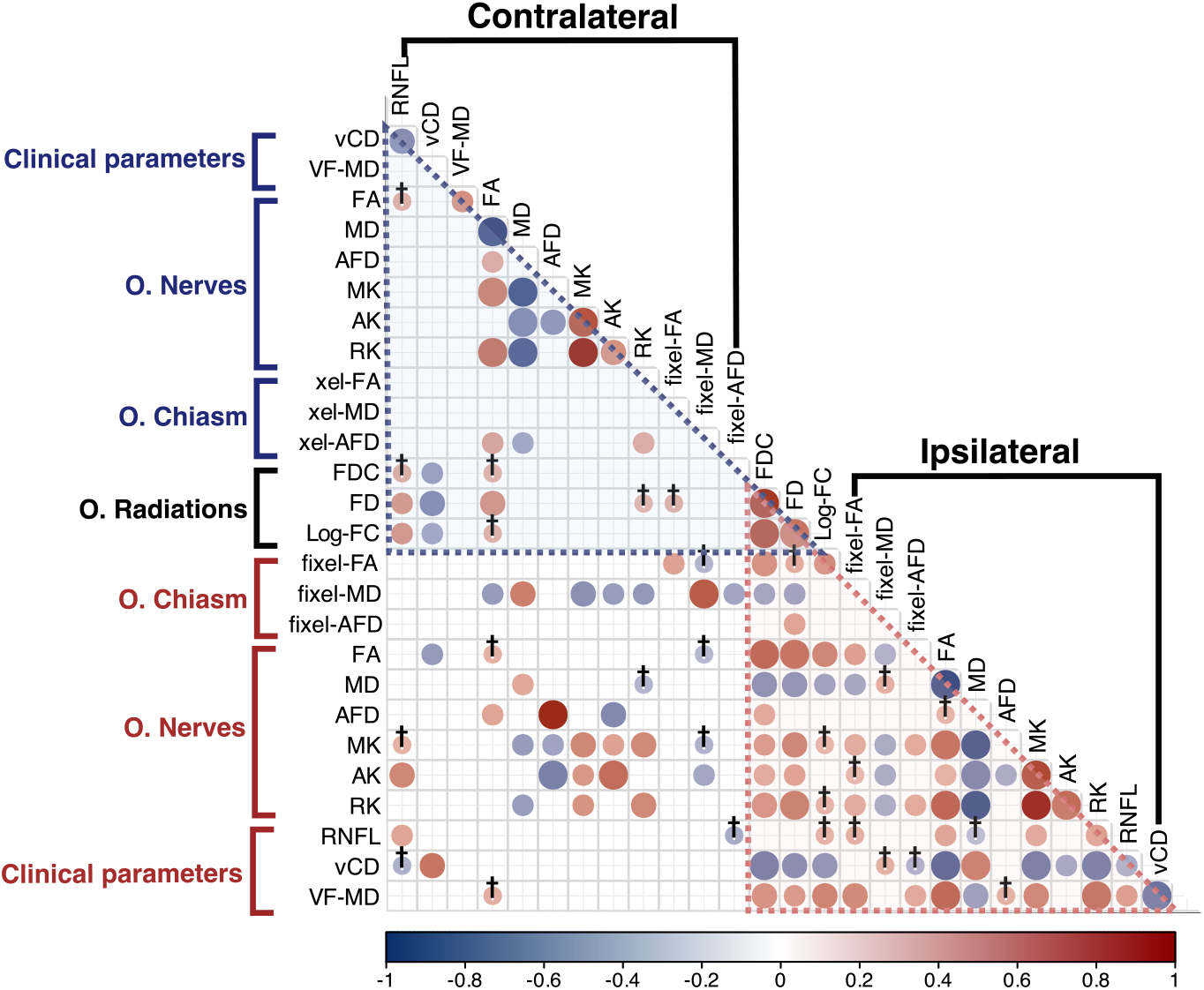
Correlation matrix of all diffusion metrics and clinical parameters organized ipsilateral and contralateral to the most affected eyes. Significant correlations are shown as circles (blue = negative, red = positive). Circle diameters reflect the significance of the correlation (p*<*0.05), and crosses indicate trends (0.05*<*p*<*0.1). The average diffusion metrics of both hemispheres is used for the optic radiations

## Acknowledgments

We thank Dr. Erick Pasaye for support during MRI acquisition. Data analysis was partially performed at the National Laboratory for advanced scientific visualization (LAVIS), with help from Luis Aguilar. We also appreciate the technical support provided by Mirelta Regalado, Leopoldo González-Santos, Juan Ortiz-Retana and Moisés Baltazar. Dr. Ricardo Ríos and Dr. Stéphanie Thebault provided useful ideas and suggestions during the course of this project. Daniela Coutiño is a doctoral student from Programa de Doctorado en Ciencias Biomédicas, Universidad Nacional Autónoma de México (UNAM) and was supported by CONAHCYT scholarship (CVU 1288997). Funding was provided by UNAM (PAPIIT IN213423) and CONAHCYT (CF-2023-I-218). The National Laboratory for MRI has received funding from CONAHCYT/SECIHTI.

